# Algebraic Shortcuts for Leave-One-Out Cross-Validation in Supervised Network Inference

**DOI:** 10.1101/242321

**Authors:** Michiel Stock, Tapio Pahikkala, Antti Airola, Willem Waegeman, Bernard De Baets

## Abstract

**Motivation:** Supervised machine learning techniques have traditionally been very successful at reconstructing biological networks, such as protein-ligand interaction, protein-protein interaction and gene regulatory networks. Recently, much emphasis has been placed on the correct evaluation of such supervised models. It is vital to distinguish between using the model to either predict new interactions in a given network or to predict interactions for a new vertex not present in the original network. Specific cross-validation schemes need to be used to assess the performance in such different prediction settings.

**Results:** We present a series of leave-one-out cross-validation shortcuts to rapidly estimate the performance of state-of-the-art kernel-based network inference techniques.

**Availability:** The machine learning techniques with the algebraic shortcuts are implemented in the RLScore software package.

## 1 Introduction

Biological systems can be understood as large collections of interacting parts, such as genes, proteins, nucleic acids and small organic molecules. Computational techniques are required to model such biological networks, since the number of possible interactions is simply too large to explore experimentally. Furthermore, high-throughput screenings are often noisy and not always reproducible [Bonetta, 2010, Prinz et al., 2011]. Supervised machine learning techniques have been successfully used for biological network inference for over a decade [Ben-Hur and Noble, 2005, Vert, 2008, Ding et al., 2013, Schrynemackers et al., 2013]. Such methods depart from an experimentally determined network, a set of observed interactions, from which a statistical model is learned. This model can subsequently be used to suggest missing interactions in the given network or to predict interactions with new vertices. Despite the fact that supervised network inference is arguably only a specific application of standard regression or classification algorithms, there are some specific challenges in correctly estimating the performance of a learned model [Park and Marcotte, 2012, Schrynemackers et al., 2013, Pahikkala et al., 2015].

The network used to build the model can be represented as an adjacency matrix, with the rows and columns representing the vertices and the matrix elements the values of the edges between the vertices. If the network is only characterized by the presence or absence of an interaction, these values are binary and the matrix is often sparse; if the interaction strength is measured, for example in the form of a binding affinity, the elements of the adjacency matrix are real-valued. In supervised network inference, one also uses a description of the vertices, typically in the form of numerical features (e.g. a molecular fingerprint) or a similarity matrix (e.g. obtained by sequence alignment). Using the example network and these vertex descriptors, a function is learned to predict the interaction value for two given vertices.

A major problem in assessing the performance of models for supervised network inference is that it is not unambiguously clear how to choose an independent test set. Following the work of [Park and Marcotte, 2012], we distinguish different prediction settings, dependent on whether one is interested in detecting new interactions between the vertices of the training network or predicting for one or two new vertices. The performance for each of those settings should be assessed accordingly. For each of the settings, we present computational shortcuts to perform suitable leave-one-out (LOO) cross-validation, allowing for extremely rapidly tuning and validating models for such settings. These shortcuts relate to a kernel-based method for network inference, namely two-step kernel ridge regression [Pahikkala et al., 2014, Romera-Paredes and Torr, 2015], a state-of-the-art method for network inference.

The first set of cross-validation schemes applies to bipartite networks, i.e. graphs for which the vertices can be subdivided in two disjoint sets and edges only occur between vertices of different sets. Examples of bipartite network inference include protein-ligand interaction prediction [Jacob and Vert, 2008, Bleakley and Yamanishi, 2009, van Laarhoven et al., 2011, Gönen, 2012, Ding et al., 2013, Li et al., 2016], mRNA-miRNA interaction prediction [Van Peer et al., 2016] and nucleic acid-protein affinity prediction [Pelossof et al., 2015]. Suppose that one wants to build a model to predict protein-ligand interaction strength. For a given protein-ligand pair, four predictions settings can be distinguished: (1) the protein and the ligand both occur in the training network, (2) only the protein occurs in the training network, (3) only the ligand occurs in the training network or (4) both vertices are new. For these four settings, we define four respective leave-one-out cross-validation settings. In Setting I, one interaction or value of the adjacency matrix is withheld at a time. In Setting R, every row of the adjacency matrix is withheld once. Similarly, in Setting C, every column is withheld one-by-one. Finally, in Setting B, every element of the adjacency matrix is withheld once and the model is trained using the adjacency matrix with both the row and column containing that element discarded. These settings are depicted in the top of Figure 1. Note that even though the toy example illustrates network inference as a binary classification task (predict presence/absence of an interaction), our settings can also be used for regression tasks such as predicting binding affinities between molecules.

**Fig. 1.**
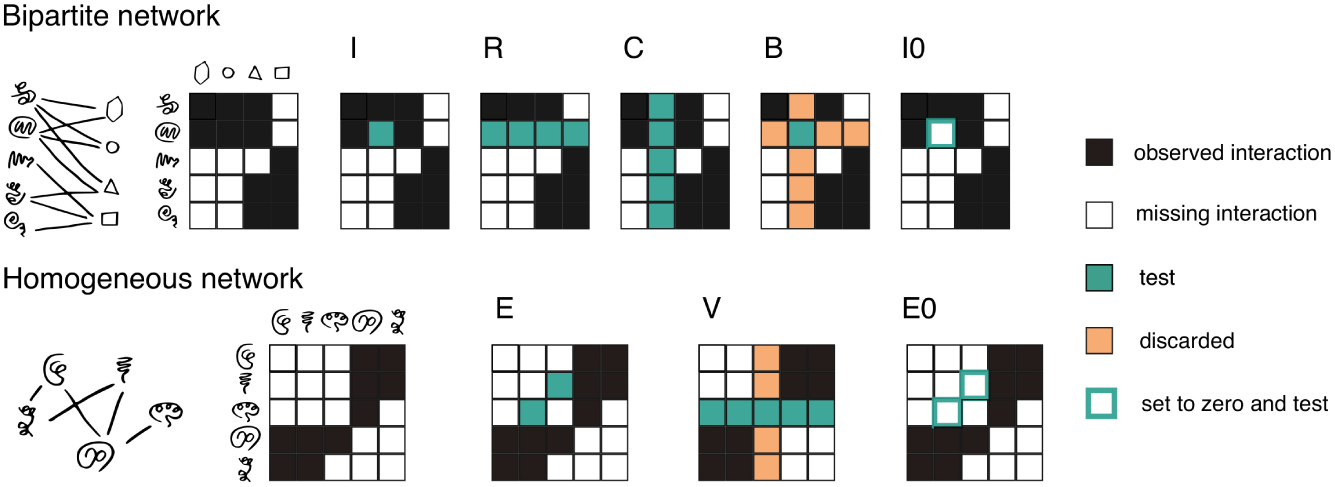
Overview of the network inference problems discussed in this work. (top) Bipartite network prediction, where the vertices are of a different type, e.g. predicting interactions between proteins and ligands. A toy network between five proteins and four ligands with the interaction network is shown. Here, we distinguish four prediction settings: I (interactions), R (rows), C (columns) and B (both). (bottom) Network prediction, where the vertices are of the same type, e.g. protein-protein interaction prediction. A toy network of interactions between five proteins is shown. Here, two prediction settings are distinguished: E (edges) and V (vertices). We also present variants of both Settings I and E: Settings I0 and E0. Here, the value of an interaction is set to zero, rather than being discarded. See main text for details.

Some slightly different prediction settings arise for homogeneous networks, i.e. networks for which the vertices are of the same kind. Inference problems for these types of networks arise for protein-protein interaction networks [Vert et al., 2007, Hamp and Rost, 2015, Liu et al., 2015, Schrynemackers et al., 2015], gene regulatory networks [Huynh-Thu et al., 2010, Marbach et al., 2012, Maetschke et al., 2014] and metabolic networks [Yamanishi et al., 2005, Geurts et al., 2007]. Homogeneous networks are represented by square adjacency matrices. In addition, the adjacency matrix is often symmetric (in case of proteinprotein interaction networks) or, more rarely, skew-symmetric (in metabolic networks, an ingoing flux is always accompanied by a negative outgoing flux of equal magnitude). To accommodate for these properties, we suggest two additional prediction settings for homogeneous networks: predicting for edges within the network or predicting how new vertices will interact with the existing network. We refer to the former setting as Setting E. Here, we remove one edge of the network at a time and predict for this edge. This edge is represented by two values in the adjacency matrix: one above (i.e. the element at position (*i,j*)) and one below the diagonal (i.e. the element at position (*j,i*)). To evaluate how new vertices interact with the network, we suggest Setting V. Here, every vertex is removed once from the network and the interaction values of the remaining vertices with this left-out vertex are predicted. Leaving out one vertex corresponds to removing the corresponding row and column of the adjacency matrix. These two cross-validation settings are depicted in the bottom part of Figure 1.

Biological networks have one more peculiarity to keep in mind: the occurrence of false negatives. When networks are constructed by experimentally determining all the pairwise interactions, e.g. an assay of kinase inhibition, all interactions are assumed to be correct within experimental error. Usually, however, biological networks are obtained by aggregating positive interactions. This means that there is often no experimental evidence for the absence of an interaction. Missing links in a network are either true negative interactions or false negative ones, i.e. the interaction between the vertices is simply not observed. Several researchers [Elkan and Noto, 2008, Cerulo et al., 2010, Schrynemackers et al., 2013, Liu et al., 2015] discussed how to train supervised models in the absence of true negative interactions. To assess whether a model can detect missing interactions within a network, we propose a small modification of Settings I and E. Rather than withholding interactions, in Setting I0, each interaction of the adjacency matrix is in turn set to zero, regardless whether there was a non-zero interaction value or not. Thus, for every element in the adjacency matrix, the value is set to zero, and a prediction is made for that element using a model trained with the modified adjacency matrix. The same principle is used in Setting E0. Both variants are also depicted in Figure 1. Here, the values of edges are in turn set to zero. Such cross-validation schemes have been used, for example, by [van Laarhoven et al., 2011]. Depending on whether one can deal with false negatives or not, Setting I, resp. Setting E, is more relevant than Setting I0, resp. E0.

The remainder of this work is structured as follows. Section 2 outlines the two-step kernel method for network inference. In Section 3 we give the computational shortcuts for cross-validation in network inference. In Section 3.1 the basic shortcuts for kernel ridge regression are presented. These are combined in Sections 3.2 and 3.3 into the shortcuts for bipartite and homogeneous networks, respectively. The shortcuts are illustrated on some benchmark biological networks in Section 4. Finally, in Section 5 we end this work with some pointers for future research.

We use boldface small cap letters for vectors, e.g. **a**, and capital letters for matrices, e.g. *A*. The *i*-th element of vector **a** is denoted by *a_i_*. We use *A_i_*. to denote the *i*-th row vector of *A*, *A_.j_* to denote the *j*-th column vector of *A* and *A_ij_* for the element at position 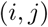 of matrix *A*. Similarly, for sets of indices 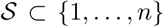 and 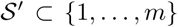, 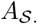, 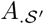, and 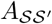 denote the submatrices of *A* in which the rows, columns or both are indexed by 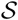 or 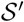.

## 2 Supervised network inference with two-step kernel ridge regression

### 2.1 Predicting bipartite networks

Supervised network inference starts from a set of labeled interactions. For bipartite networks, one wants to predict an interaction between vertices of two different kinds, e.g. between proteins and ligands or between miRNAs and mRNAs. There are thus two object spaces, 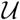 and 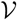. Suppose the training set contains information on the subsets 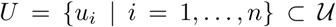 and 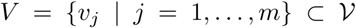 of both types of objects. There is exactly one observed label *Y_ij_* for every pair 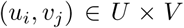, stored in the *n* × *m* adjacency matrix *Y*. These labels can either be binary, indicating the presence or absence of an interaction, or real-valued, indicating for example the strength of an interaction or a kinetic parameter.

The goal of supervised network inference is to learn a predictive model 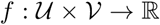 that takes a pair of vertices as input and returns a value that either estimates the interaction label or that can be interpreted as a score indicating the confidence in the occurrence of an interaction. In kernel methods, such a function is based on a feature description of both vertices by means of kernel functions. Kernels are symmetric and positive-definite functions that quantify the similarity between vertices. They are very popular in bioinformatics because they can be used to implicitly represent structured objects such as sequences, trees and graphs [Shawe-Taylor and Cristianini, 2004]. Let 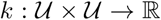 be the kernel function associated with vertices in 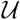 and, likewise, 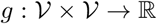 be the kernel function associated with vertices in 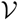. The model to be learned is of the form

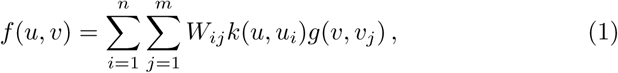

with *W* = [*W_ij_*] an *n* × *m* matrix of weights. Models of this type are commonly used for biological network inference, e.g. [Ben-Hur and Noble, 2005, Vert et al., 2007, Jacob and Vert, 2008, van Laarhoven et al., 2011, Pahikkala et al., 2015, Pelossof et al., 2015]. They arise naturally when using the Kronecker kernel in a kernel-based learning algorithm such as a support vector machine or kernel ridge regression. Such pairwise kernel methods achieve state-of-the-art performance for many network inference tasks.

In this work, we will focus on the two-step kernel ridge regression method for fitting the model of Eq. (1). This method was independently proposed by [Pahikkala et al., 2014] and [Romera-Paredes and Torr, 2015], though similar methods have been proposed earlier in structural equation modeling [Bollen, 1996, Jung, 2013]. We introduce the Gram matrices:

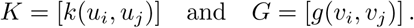

The parameters for two-step kernel ridge regression are obtained as

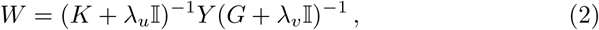

with I the identity matrix and λ*_u_* and λ*_v_* two regularization parameters that have to be tuned. Equation (2) can be motivated in several ways. Firstly, it is obtained when, as the name suggests, two successive kernel ridge regression steps are executed: once to generalise to new objects in 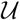 and once to generalise to new objects in 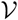. Secondly, Eq. (2) can also be seen as solving the inverse problem of finding the parameters *W* of model (1) to explain the observed labels *Y*. The regularization is then required to make this inverse problem well-posed, such that tiny fluctuations in the labels do not result in huge changes in model behavior. Regardless of how two-step kernel ridge regression is derived, it is shown theoretically to be a well-founded method [Stock et al., 2017].

The associated matrix of predictions, 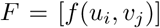, can easily be computed as

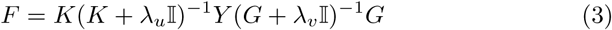

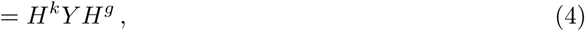

where 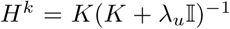 and 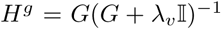 are further on referred to as the hat matrices. Note that given the eigenvalue decompositions of the Gram matrices, i.e.

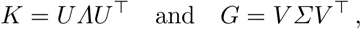

the hat matrices can easily be obtained as

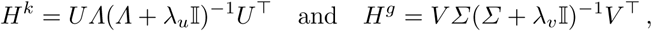

for any values of the regularization parameters. Note that in the above computations one only has to invert diagonal matrices, which can be done by simply inverting the element on the diagonal. The eigenvalue decompositions can be computed with a time complexity of 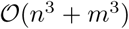. Given this decomposition, the model weights and the predictions can be obtained for any value of the regularization parameters by matrix multiplication.

### 2.2 Predicting homogeneous networks

A special, yet important case occurs when interactions between objects of the *same* kind are predicted, e.g. protein-protein interaction networks, metabolic networks or gene regulatory networks. Here, there is only one set of objects 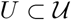 and the adjacency matrix *Y* is square. The equations hence simplify to

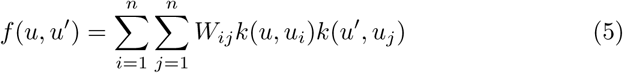

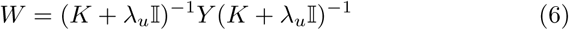

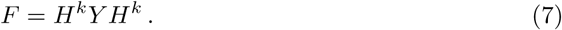

In some cases, some special structure can be imposed on the adjacency matrix. When the label of every pair (*u*, *u*′) is the same as the label of (*u*′, *u*), this is called a *symmetric* interaction. Such relations occur for example in protein-protein interaction networks. For symmetric relations, we expect that *f*(*u*, *u*′) = *f*(*u*′, *u*) for any 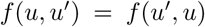. Likewise, *skew-symmetric* prediction functions satisfy that *f*(*u*, *u*′) = − *f*(*u*′, *u*) for any 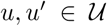. Skew-symmetric functions are of relevance when modelling gene regulatory networks and flows in a metabolic network.

Whenever *Y* is symmetric, the learned prediction function will also be symmetric, i.e. *W* = *W*^⊤^. This can be seen from Eq. (6). The same holds for skew-symmetric adjacency matrices, where a skew-symmetric adjacency training network will lead to parameters that satisfy *W* = − *W*^⊤^.

## 3 Shortcuts for leave-one-out cross-validation

Two-step kernel ridge regression is, as implied by its name, merely performing kernel ridge regression twice. As such, traditional leave-one-out crossvalidation shortcuts can be used as building blocks towards the shortcuts for the more complex network cross-validation schemes. In Section 3.1, the shortcuts for kernel ridge regression are presented. Sections 3.2 and 3.3 present the shortcuts for the network cross-validation settings depicted in Figure 1. The formal derivations of these shortcuts are presented in the supplementary materials.

### 3.1 Basic shortcuts for leave-one-out cross-validation

The traditional leave-one-out shortcuts can be applied for any model that minimizes a squared loss and where a vector of labels *Y_i_*. (here rows from the adjacency matrix) and the corresponding vector of predictions *F*_i._ are linked through a hat matrix [Wahba, 1990]:

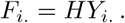

Crucially, the hat matrix depends *only* on the feature descriptions of the instances and *not* on the labels. Popular methods such as (kernel) ridge regression, splines, spectral regularization methods and extreme learning machines fall into this category. Figure 2 shows a toy regression problem to illustrate the different shortcuts.

**Fig. 2.**
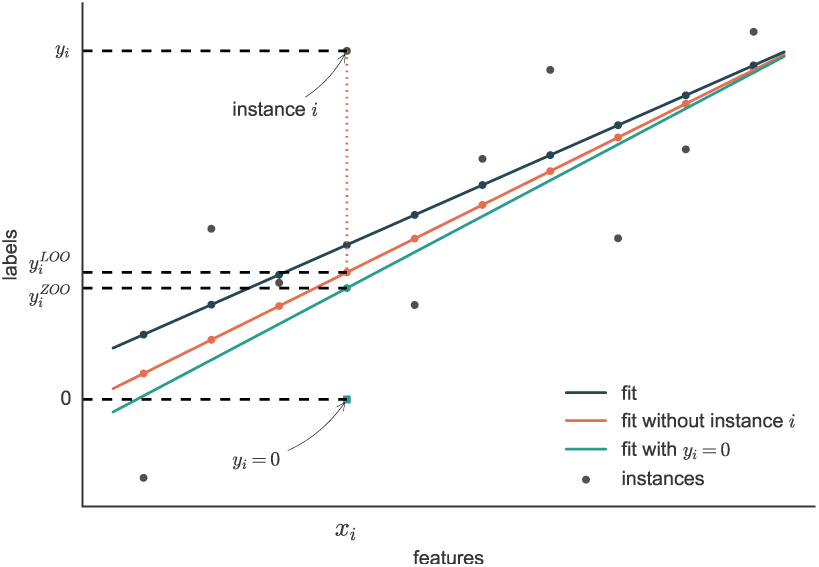
Toy regression problem with 10 instances. The *i*-th instance has label *y_i_* and a single feature *x_i_*. The blue line is a ridge regression model fit on the data, the red line is the fit without the *i*-th instance and the green line is the fit in which the label of the *i*-th instance is set to zero.

The following theorem gives the shortcut to compute the corresponding predictions for an adjacency matrix with the rows indexed by S removed.

#### Theorem 1

*For an n* × *m matrix of labels Y and an n × n hat matrix H, the elements indexed by the set 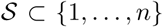 of the leave-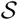-out split are given by the* 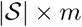 *matrix*

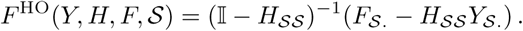

The proof of the theorem is given in the supplementary materials. The main idea behind is that if one replaces the rows indexed by 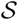 by the corresponding rows of *F^HO^*(*Y*, *H*, *F*, *S*) in the label matrix, the corresponding predictions using a model trained using these labels again yield *F^HO^*(*Y*, *H*, *F*, *S*). This is illustrated in Figure 2, in which the LOO prediction for instance i coincides with the corresponding prediction of the model fitted without the *i*-th instance.

Rather than removing an instance, one can also opt for setting the value to zero. Such setting is relevant to assess whether the model can detect false negatives, i.e. whether a zero should be a one in the adjacency matrix. We use the term zero-one-out (ZOO) for the scheme in which models are fitted using data where the labels are set to zero one-by-one. The following theorem gives a computational shortcut to compute the predictions for an adjacency matrix with an arbitrary number of rows set to zero.

#### Theorem 2

*For an n* × *m matrix of labels Y and an n × n hat matrix H, if the elements indexed by the set* 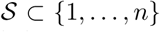 *are set to zero, the corresponding predictions are given by the* 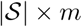 *matrix*

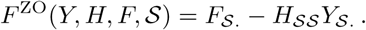

### 3.2 Leave-one-out in bipartite networks

Table 1 shows the shortcuts for the different cross-validation settings for bipartite networks. The shortcuts for Settings I and I0 can easily be deduced by transforming the network prediction problem into a standard prediction problem using the vec operator, which stacks the columns of a matrix into a vector. As such, Eq. (3) becomes

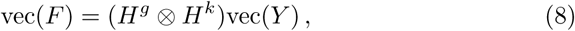

with ⊗ the Kronecker product. We have made use of the identity *vec*(*AXB*) = (*B*^⊤^ ⊗ *A*)vec(X), which holds for any conformable matrices *A*, *B* and *X*. Theorems 1 and 2 then yield the shortcuts for Settings I and I0. The shortcuts for the Settings R, C and B can be obtained by considering the two steps of two-step kernel ridge regression separately and applying Theorem 1 on the rows, columns or both, respectively.

**Table 1.**
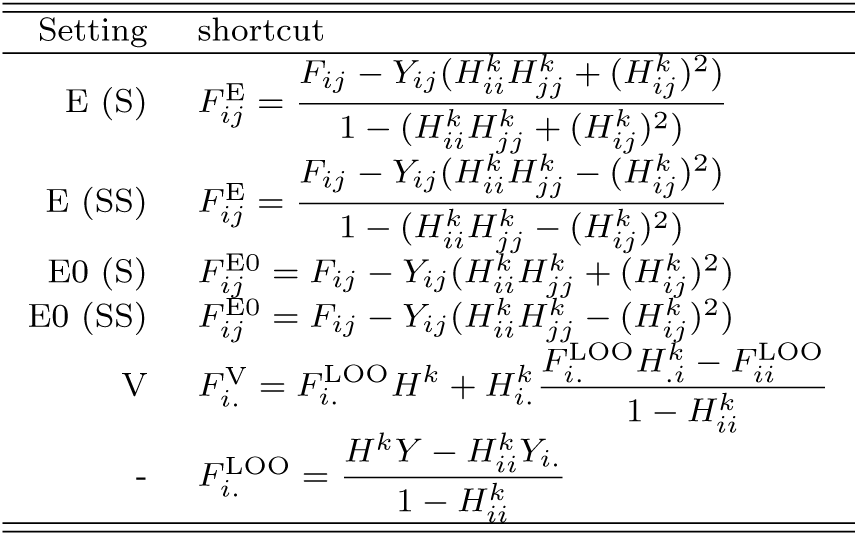
The different shortcuts for bipartite networks. As shown in Figure 1, the settings I (interactions), R (rows), C (columns) and B (both) are considered. The variation of Setting I, Setting I0, is when a single element at position (*i,j*) is set to zero, rather than being withheld. For bipartite networks *F* = *H^k^YH^g^*.

The shortcuts for bipartite networks have an approximate time complexity of 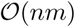, given that the hat matrices and the prediction matrix have been precomputed. In the case of Settings R, C and B, the matrices *H^k^Y* and *YH^g^*, intermediate results towards computing *F*, should also be kept in cache to reach this time complexity.

### 3.3 Leave-one-pair-out in homogeneous networks

Homogeneous networks impose additional dependencies in the adjacency matrices. To derive the shortcuts for leaving out edges, Eq. (7) can also be stated in vector form as Eq. (8). By leaving out the values at positions (*i,j*) and (*j, i*) and using the properties of symmetry and skew-symmetry, the shortcuts for Setting E can be derived. The shortcut for Setting V can be obtained by applying the shortcut of Theorem 1 twice. The shortcuts for homogeneous networks are summarized in Table 2. Given that *F* and *H^k^* are precomputed, these leave-out-out adjacency matrices can be computed with a time complexity of 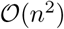, i.e. constant time complexity for each element.

**Table 2.**
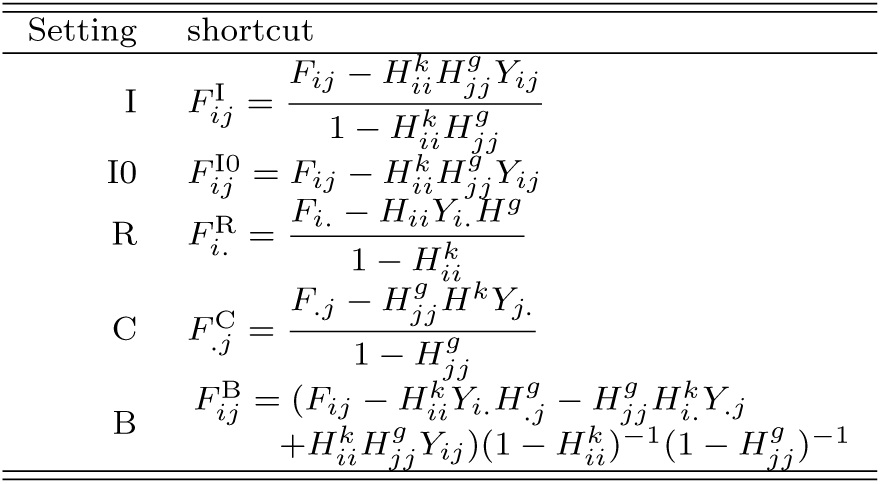
The different shortcuts for homogeneous networks. As shown in Figure 1, Settings E (edges) and V (vertices) are considered. The shortcuts for Setting E can be used for symmetric (S) and skew-symmetric (SS) matrices. The variation of Setting E, Setting E0, is when the label of the edge from vertex *i* to vertex *j* is set to zero, rather than being withheld. For homogeneous networks *F* = *H^k^YH^k^*.

## 4 Experiments

In the experiments we will illustrate the speed of the leave-one-out shortcuts provided in this work. To this end, we use four benchmark datasets of bipartite protein-ligand interaction networks collected by [Yamanishi et al., 2008]^1^ and one homogeneous protein-protein interaction dataset from [Vert et al., 2007]. These datasets are described in Table 3. In Section 4.1 we compare the shortcuts for the bipartite networks with Kronecker kernel ridge regression, whereas in Section 4.2 we study the scalability of the shortcuts for homogeneous networks.

### 4.1 Bipartite networks

In this section we illustrate the leave-one-out shortcuts when choosing the optimal regularization parameters for bipartite network prediction. We use the four protein-ligand datasets of [Yamanishi et al., 2008]. Each dataset concerns a different class of protein targets: enzymes (*e*), G protein-coupled receptors (*gpcr*), ion channels (*ic*) and nuclear receptors (*nr*). The interactions are given in the form of a binary adjacency matrix. Both the drugs and targets come along with a respective similarity matrix. For the drugs, common substructures are calculated using a graph alignment algorithm. The Jaccard similarity measure is used to obtain a drug similarity based on these substructures. The similarity matrix of the targets is a normalized version of the scores obtained by Smith-Waterman alignment [Smith and Waterman, 1981].

We compare two-step kernel ridge regression with Kronecker kernel ridge regression. Kronecker kernel ridge regression has only one regularization parameter *λ*, whereas two-step kernel ridge regression has two regularization parameters, *λ_u_* and *λ_v_*. Hence, if the optimal regularization parameter is sought using grid search of d values, two-step kernel ridge regression requires *d*^2^ computations of the performance rather than d for Kronecker ridge regression. For this reason, we also consider a version of two-step kernel ridge regression where *λ_u_* = *λ_v_* = *λ*, such that also a one-dimensional grid of hyperparameters has to be explored. This trade-off results in a faster tuning at the cost of potentially obtaining a slightly inferior model. In the experiments, the regularization parameters *λ*, *λ_u_* and *λ_v_* are selected from {10^−7^,10^−6^,…, 10^5^,10^6^}.

For each dataset, we perform the leave-one-out cross-validation for Settings I, R, C and B for the different values of the regularization parameters. The prediction matrices obtained using cross-validation are evaluated using micro-wise AUC (i.e. AUC computed over the complete adjacency matrix). For Setting I, a computational shortcut is available for all models. For the remaining settings, only two-step kernel ridge regression has the respective shortcuts. Hence, in the case of Kronecker kernel ridge regression, at least one eigenvalue decomposition has to be performed when withholding a protein, ligand or both.

**Table 3.** Overview of the different biological networks discussed in this work. We use four bipartite networks, enzymes (*e*), G protein-coupled receptors (*gpcr*), ion channels (*ic*) and nuclear receptors (*nr*), and one homogeneous protein-protein interaction (*ppi*) network.

| dataset | *e* | *gpcr* | *ic* | *nr* | *ppi* |
| --- | --- | --- | --- | --- | --- |
| # targets | 664 | 95 | 204 | 26 | 769 |
| # drugs | 445 | 223 | 210 | 54 | - |
| fraction of interactions (%) | 0.99 | 3.00 | 3.45 | 6.41 | 1.25 |
| median degree targets | 2 | 3 | 5 | 3 | 7 |
| median degree drugs | 2 | 2 | 3 | 1 | - |

Table 4 shows the best performances for both methods, as well as the running times for performing the complete cross-validation and hyperparameter grid search. For the different settings and datasets, we can observe that both methods have a similar performance, with two-step kernel ridge regression often slightly outperforming Kronecker kernel ridge regression. For all methods, Setting B is the hardest and Setting I the easiest, as expected.

**Table 4.**
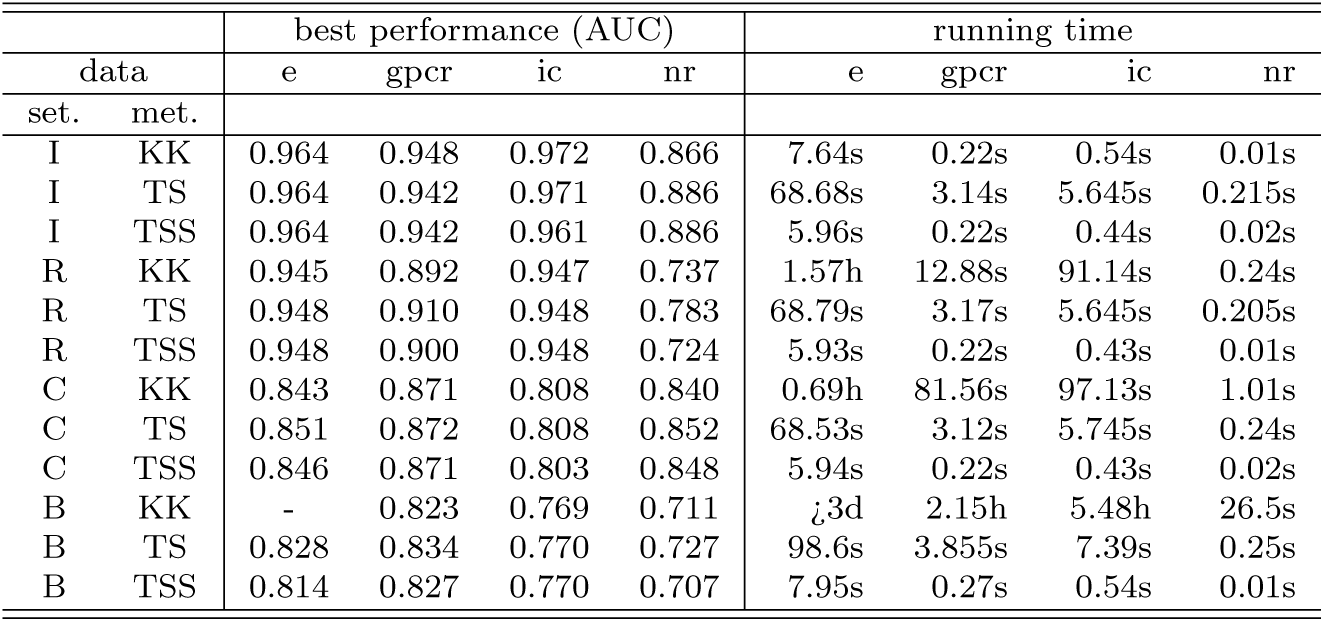
Overview of the performance and running time using Kronecker kernel ridge regression (KK), two-step kernel ridge regression (TS) and the two-step method with a single regularization parameter (TSS) for the different drug-target datasets and cross-validation settings. One experiment could not be completed in less than three days of running time. See main text for details. All experiments were performed using a basic Numpy implementation of the models and cross-validation shortcuts. All experiments were run on an AMD Opteron server (2500.159 MHz).

When comparing the running times for model selection, we can observe the computational advantage of using the shortcuts. For Setting I, both Kronecker and two-step kernel ridge regression have a holdout shortcut, hence both are fast. Kronecker kernel ridge regression has to iterate over a set of fifteen regularization values, while two-step kernel ridge regression has to search a grid of 15 × 15 regularization parameters, making the latter slower. Both methods are very fast in practice, however, since the main bottleneck is computing the eigenvalue decomposition of the two Gram matrices. For Settings R and C, there is only an efficient algorithm for two-step kernel ridge regression. For datasets larger than dataset *nr*, two-step kernel ridge regression is much faster than Kronecker kernel ridge regression. For Setting B two-step kernel ridge regression is several magnitudes faster compared to Kronecker kernel ridge regression. For the latter method it was not even possible to perform this cross-validation for the *e* dataset within three days.

In the supplementary materials, we show how the performance of the different methods changes with the different regularization parameter(s). Two conclusions can be drawn from these experiments. Firstly, the performance is quite sensitive to the chosen regularization parameters. Secondly, the optimal regularization parameters are quite different, depending on the prediction setting that is evaluated. This illustrates the importance of the four crossvalidation settings discussed in this work. Using the shortcuts, the best model for the given task can be obtained.

### 4.2 Homogeneous networks

In this section, we explore how the computing time scales for performing crossvalidation in homogeneous networks. We again use the four protein-ligand networks of the previous settings, though they are turned into eight square matrices by either multiplying a label matrix with its transpose or multiplying the transpose of an adjacency matrix with the adjacency matrix itself. These represent the number of molecules two proteins or ligands have in common as binding partner. The goal here is to predict for a given molecule how many indirect interactions there are with another molecule of the same type. Though this setting is arguably somewhat artificial, it is well suited to demonstrate our shortcuts. The values of the new label matrices are variance-stabilized by means of a square root transformation.

In addition to the protein-ligand networks, we also use the protein-protein interaction (*ppi*) network of [Vert et al., 2007]. In this work, the proteins are described using seven different Gram matrices, encoding information on the location, expression, phylogeny, etc. Following the original paper, we summed these kernel matrices as this resulted in the best performing model.

For the different datasets, we measured the time to train a model (i.e. compute the hat matrix *H*), to make a prediction (i.e. compute the prediction matrix *F* given *H*) and to compute the leave-one-out matrices for Settings E and V using our shortcuts. Every computation was done for values of λ in {10^−5^, 10^−4^,…, 10^4^,10} and we computed the average time over the 11 experiments.

Figure 3 shows how the computing time scales with the number of vertices in the network. Computing the hat matrix is dominated by the eigenvalue decomposition of the hat matrix and takes the most time. Computing the prediction matrix given the hat matrix takes the least time as this only involves multiplying three matrices. The time to compute the cross-validation matrices takes an intermediate amount of time, though for larger networks these times seem to converge asymptotically to the prediction time. For reference, we added the times it would take to naively compute the cross-validation matrices. For Setting E, resp. Setting V, we took 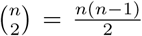, resp. n, times the training time plus one time the prediction time. Again, our shortcuts are many orders of magnitude faster than computing the cross-validation naively. We refer to the supplementary materials for an overview of the performance of the different models.

**Fig. 3.**
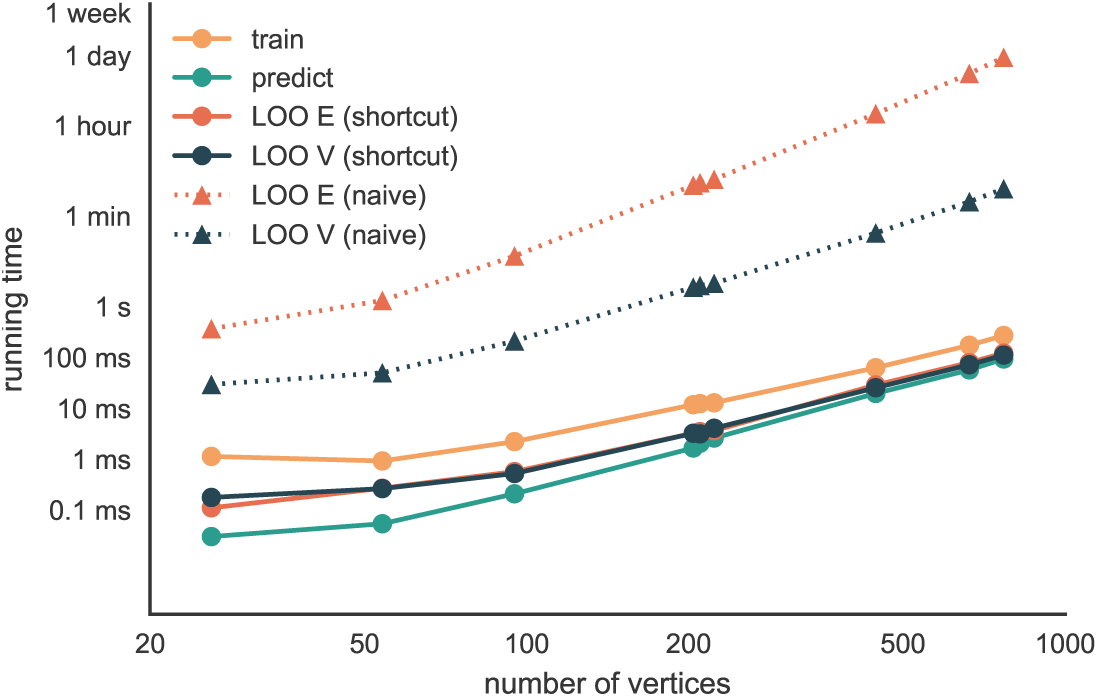
Time for training, predicting and leave-one-out computation in several homogeneous networks. For the training time, the time to construct the hat matrix is taken. Constructing the leave-one-out matrices for Settings E and V using the provided shortcuts takes less time than training, but more than for computing the prediction matrix. The time for computing leave-one-out matrices naively is estimated from the training and prediction times. Experiments performed on a MacBook Pro 2.5 GHz Intel Core i5.

## 5 Discussion and conclusion

In this work we have presented a series of algebraic shortcuts for leave-one-out cross-validation for the biological network inference problem. These shortcuts are a valuable tool for selecting the best model and for accurately estimating the model performance on new data. The shortcuts apply to a simple, though powerful network inference model, two-step kernel kernel ridge regression. Kernel methods are generally liked by the computational biology community, both because they are strong learners and because prior knowledge can naturally be assimilated. Given the eigenvalue decomposition of the Gram matrices of the vertices, leave-one-out cross-validation can be performed for any of the discussed settings and any values of the regularization parameters in roughly the time needed to make a prediction of the original adjacency matrix. Leave-one-out cross-validation provides nearly unbiased, though sometimes high variance estimates of the generalization error of a model. This is because all the models are trained using largely the same data. Our shortcuts can easily be extended to leaving out larger blocks of the adjacency matrix using Theorem 1.

Rather than developing new machine learning techniques to learn from the given small interaction networks, we believe that the largest progress in biological network inference will be made by using larger datasets and constructing better feature representations for the vertices. Randomized algorithms allow for approximating the decomposition of huge Gram matrices [Gittens and Mahoney, 2013] or constructing a nonlinear feature description directly [Huang et al., 2015]. Recent advances in convolutional neural networks have resulted in intriguing ways to generate representations for molecules [Duvenaud et al., 2015], proteins [Jo et al., 2015] and nucleic acids [Alipanahi et al., 2015]. In such cases, one would prefer to work in the primal form and the hat matrix is hence computed as

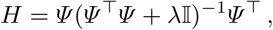

with *Ψ* a feature matrix. We are convinced that the framework discussed in this work will remain relevant in light of these exciting developments.

## Footnotes

1 Available at http://web.kuicr.kyoto-u.ac.jp/supp/yoshi/drugtarget

